# Short paired-end reads trump long single-end reads for expression analysis

**DOI:** 10.1101/777409

**Authors:** Adam H. Freedman, John M. Gaspar, Timothy B. Sackton

**Affiliations:** Faculty of Arts and Sciences Informatics Group, Harvard University, Cambridge, Massachusetts 02138

## Abstract

**Background:** Typical experimental design advice for expression analyses using RNA-seq generally assumes that single-end reads provide robust gene-level expression estimates in a cost-effective manner, and that the additional benefits obtained from paired-end sequencing are not worth the additional cost. However, in many cases (e.g., with Illumina NextSeq and NovaSeq instruments), shorter paired-end reads and longer single-end reads can be generated for the same cost, and it is not obvious which strategy should be preferred. Using publicly available data, we test whether short-paired end reads can achieve more robust expression estimates and differential expression results than single-end reads of approximately the same total number of sequenced bases.

**Results:** At both the transcript and gene levels, 2×40 paired-end reads unequivocally provide expression estimates that are more highly correlated with 2×125 than 1×75 reads; in nearly all cases, those correlations are also greater than for 1×125, despite the greater total number of sequenced bases for the latter. Across an array of metrics, differential expression tests based upon 2×40 consistently outperform those using 1×75.

**Conclusion:** Researchers seeking a cost-effective approach for gene-level expression analysis should prefer short paired-end reads over a longer single-end strategy. Short paired-end reads will also give reasonably robust expression estimates and differential expression results at the isoform level.

## BACKGROUND

For over a decade, RNA-seq has empowered gene expression analysis. This has led to fundamental advances in our understanding of diverse phenomena, including the evolution of gene regulation across species (1), alternative splicing (2,3), relative importance of cis vs-trans regulation (4,5), genetic underpinnings of heritable disease (6,7), and the genetic architecture of phenotypes in natural populations (8,9). While the cost of sequencing has steadily decreased, making expression analyses feasible even on modest research budgets, maximizing cost efficiency remains high priority. Cost effectiveness is particularly important as studies scale up to hundreds or even thousands of samples. Maximizing performance given a particular study design has already received much attention, with investigations of optimal strategies for quality trimming (10), the effects of quality trimming on expression estimates and differential expression testing (11), optimal sequencing depth (12,13), importance of biological replicates (12,14,15), and the relative performance of expression and differential expression tools and pipelines (16–20). Less attention has focused on how to improve performance given a fixed number of biological replicates and total sequenced bases.

While the typical length of sequenced reads has increased as costs have decreased, when an annotated genome is available it remains standard procedure to sequence single end reads to estimate gene expression, typically around 75 base pairs. Yet, this strategy does not leverage information provided by the full length of the library fragments, particularly the greater specificity in read mapping (or pseudo-alignment) that would be provided by sequencing from both ends of a fragment. Given a fixed cost per base for the Illumina NextSeq, HiSeq X Ten, and NovaSeq instruments, paired-end 75bp reads would be twice as expensive as single-end 75bp sequencing, assuming an equivalent number of sequenced fragments. However, a sizeable fraction of the long-range information provided by paired-end sequencing should still be retained by sequencing shorter paired-end reads. In this study, we evaluate whether that intuition is correct, by assessing whether expression estimates and differential expression results obtained with 2×40 paired end sequencing are more consistent with 2×125 paired end sequencing than either 1×75 or 1×125 single-end sequencing. Our particular focus is on 1×75, as the cost of 1×75 and 2×40 are the same per read when sequencing using the Illumina NextSeq (75 cycle kit), a common instrument for RNA-seq studies. Given the fixed cost per base for the HiSeq X Ten and NovaSeq instruments, there are analogous choices to make between single-end vs. paired-end sequencing. For example, for the 100bp NovaSeq kit, a sequencing strategy trade-off exists between sequencing 2×50 vs. 1×100 reads. We conduct these analyses with publicly available Illumina sequence data across 12 different SRA (Additional file 1: Table S1). accessions encompassing diverse experimental designs and multiple model organisms with high quality annotated genomes.

## RESULTS AND DISCUSSION

Our approach to evaluating the performance of short paired-end reads vs. longer single-end ones is to compare expression estimates and differential expression results derived from these strategies to a truth set. Because we cannot know the “truth”, we assume that its closest approximation are results obtained from long, paired-end reads. Thus, we define our gold standard as the results obtained from 2×125 paired-end reads. We then trim these data down to two data sets with read lengths of 40 and 75 base pairs, and calculate Spearman’s rank correlations between transcripts-per-million (TPM) estimates based upon 2×40, 1×75 and 1×125 with the 2×125 gold standard. We present these correlations in two different ways. First, we examine the distributions of those correlation coefficients across alternative sequencing strategies. Second, we plot the correlations obtained using 2×40 over those obtained with 1×75 and 1×125.

Kallisto transcript-level TPM estimates made with 2×40 are always more highly correlated with those made with the 2×125 gold standard than those made with data trimmed to 1×75 reads (Fig. 1a,c), and perform better than estimates made with data trimmed to 1×125 for all but a few samples from one SRA accession (Fig. 1a,c,e). A similar performance advantage for the 2×40 strategy compared to the 1×75 strategy is observed at the gene level (Fig. 1 b,d,f), although there is one accession where 1×125 performs better than 2×40.

**Fig. 1.**
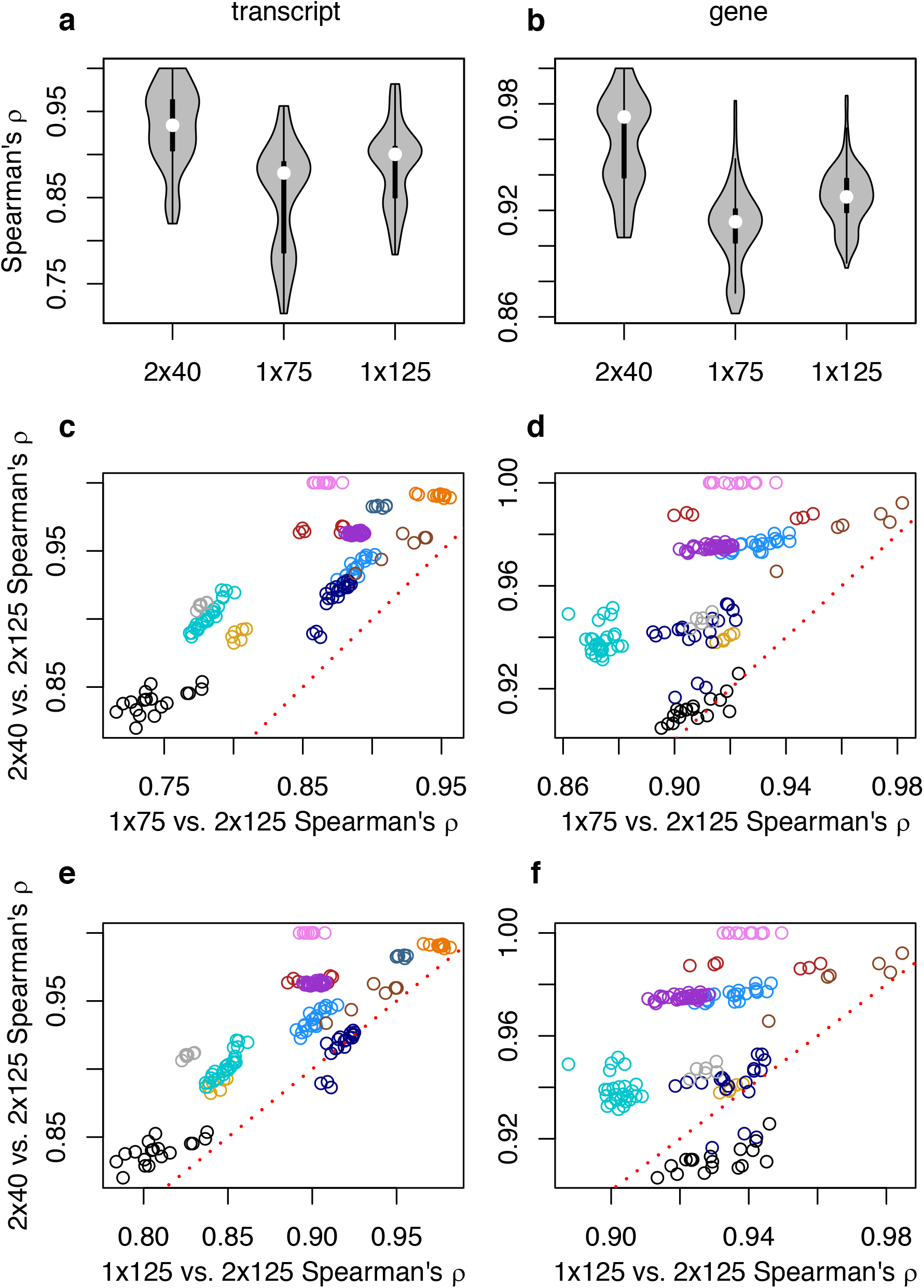
Spearman’s rank correlations for kallisto-derived transcripts per million (TPM) between the gold standard paired-end 2×125 strategy and alternative strategies. Violin plots of **(a)** transcript and **(b)** gene-level inference. Comparison of correlations with 2×125 between 2×40 and 1×75 for **(c)** transcript and **(d)** genes, and between 2×40 and 1×125 for **(e)** transcript and **(f)** genes. For **c-**f, points above the red dotted line are samples where estimates of expression from 2×40 is more highly correlated with the gold standard than the contrasted single-end strategy.

RSEM-based TPM estimates show an overall pattern generally consistent with that observed with kallisto (Fig. 2). A handful of samples for which 2×40 performed less well than both 1×75 and 1×125 had low bowtie2 alignment rates to the reference transcripts (for the 2×125 libraries, overall alignment rates of 37.1-51.0%), suggesting that library quality issues were responsible, but raising a broader issue concerning whether variation in alignment rates between strategies could explain the relative performance of 2×40 to 1×75 and 1×125.

**Fig. 2.**
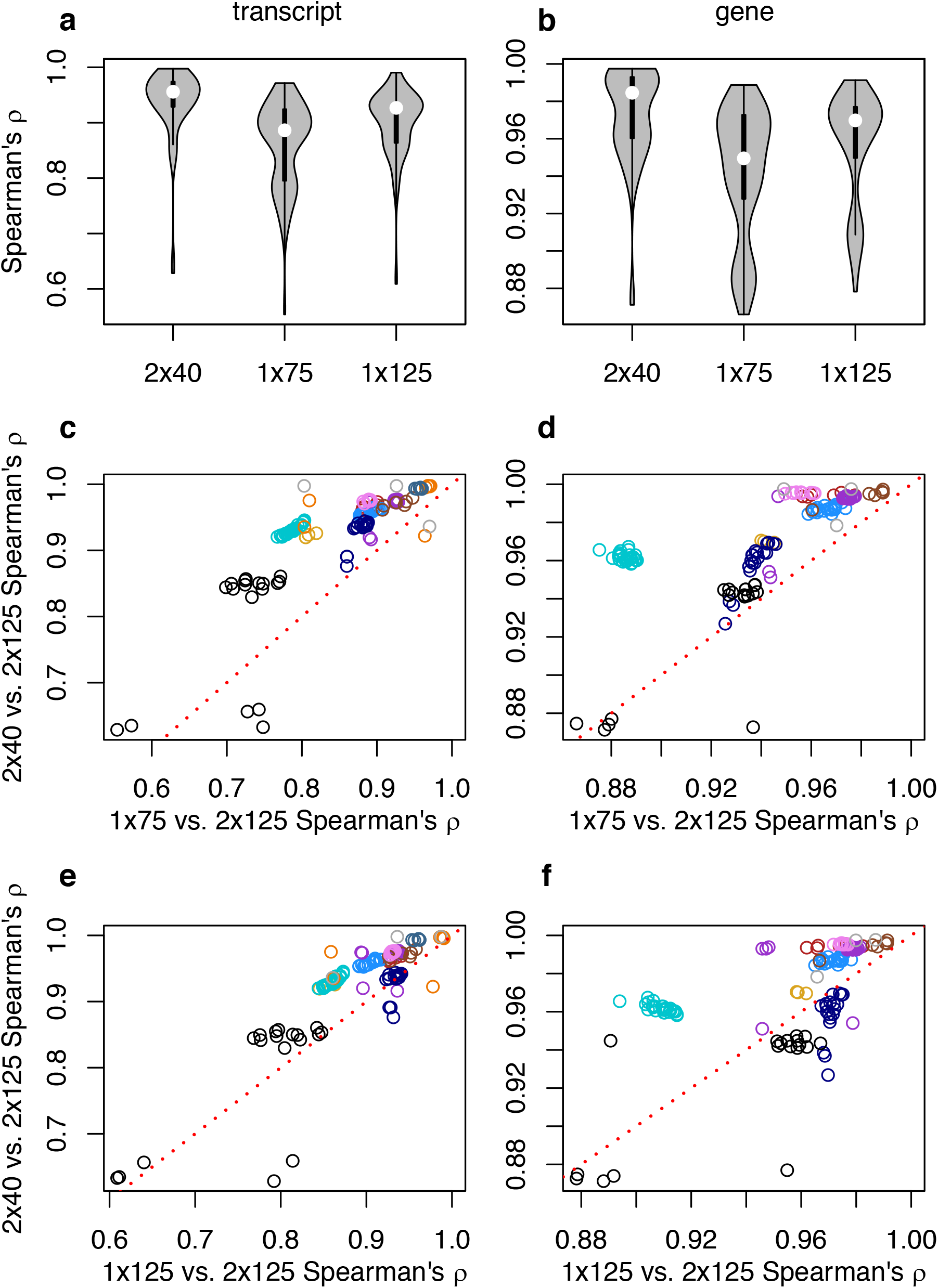
Spearman’s rank correlations for RSEM-derived transcripts per million (TPM) between the gold standard paired-end 2×125 strategy and alternative strategies. Violin plots of **(a)** transcript and **(b)** gene-level inference. Comparison of correlations with 2×125 between 2×40 and 1×75 for **(c)** transcript and **(d)** genes, and between 2×40 and 1×125 for **(e)** transcript and **(f)** genes. For **c-**f, points above the red dotted line are samples where estimates of expression from 2×40 is more highly correlated with the gold standard than the contrasted single-end strategy.

Kallisto does not conduct formal sequence alignment, so for each sample-strategy combination we summed the kallisto transcript-level counts as a proxy for the number of informative reads used in estimating expression. The summed kallisto counts for 2×40 have a weak correlation with performance (with respect to TPM) relative to 1×75 the transcript level, and a moderate correlation at the gene level (Fig 3a,b); the relative differences in counts between 2×40 and 1×75 appear uncorrelated with the relative performance of 2×40 and 1×75 (Fig. 3c,d). A similar pattern is observed with respect to 1×125 (Additional file 2: Fig. S1). For RSEM-derived expression estimates, we found no impact of the percent of reads uniquely aligning on the relative performance of 2×40, but at the gene level found that relative differences in overall alignment rate had some effect on the robustness of TPM estimates of 2×40 relative to both 1×75 and 1×125 (Additional file 3: Fig. S2,S3). For both kallisto and RSEM, while there are no obvious taxonomic effects there are clear experiment-specific effects, manifested as differences in relative 2×40 performance among accessions derived from the same species. Similar to the above results based upon expression correlations, looking across all accessions, for most samples root-mean-square error of expression estimates based upon 1×75 and 1×125 was greater than for 2×40 (Fig. 4).

**Fig. 3.**
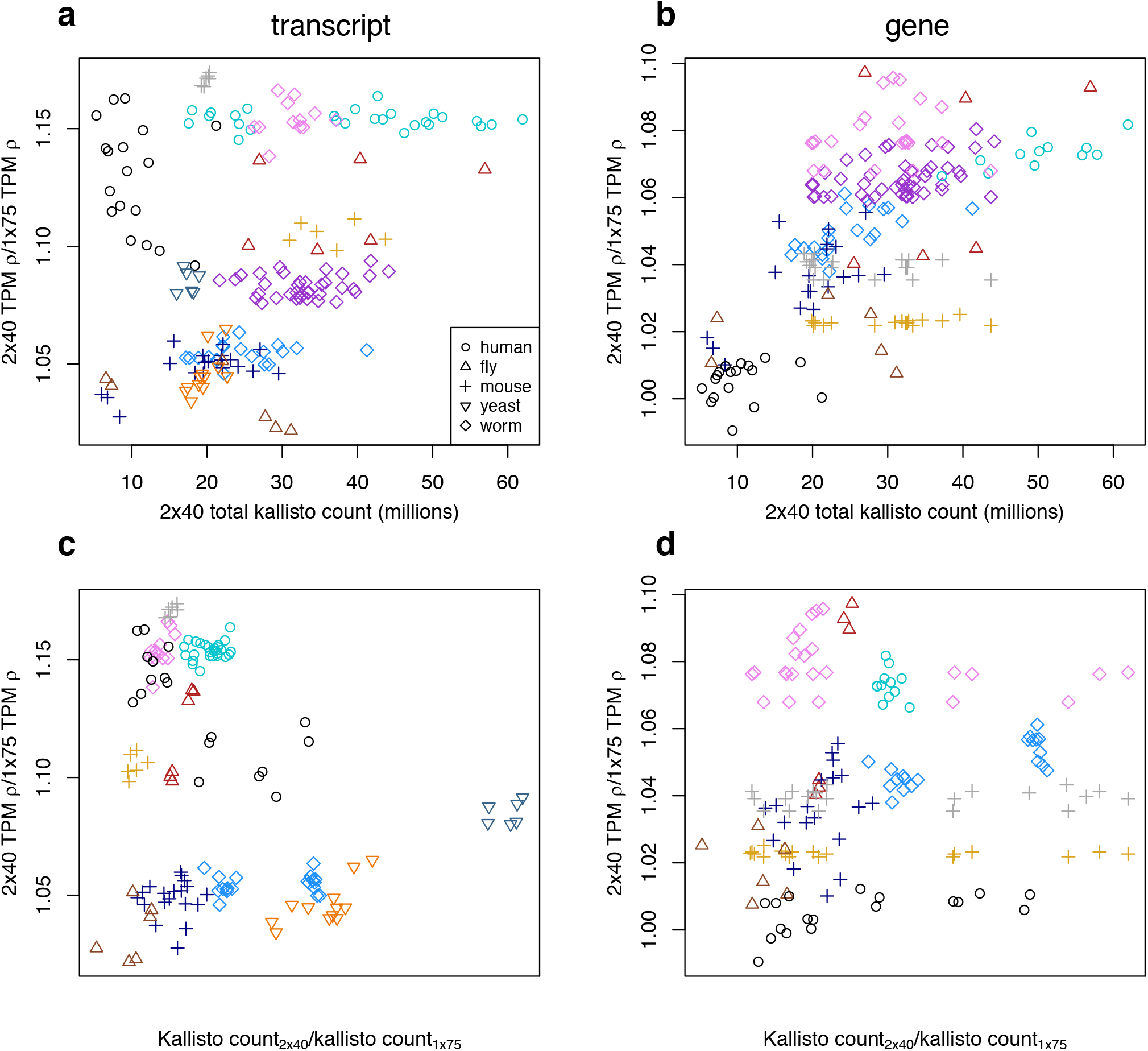
Effects of kallisto pseudo-alignment efficiency on differences in TPM estimate Spearman’s rank correlations between 2×40 vs. 1×75. Efficiency of 2×40 is presented as the total kallisto count for 2×40 for **(a)** transcripts and **(b)** genes, and contrasted with the efficiency of 1×75 in the form of count ratios for **(c)** transcripts and **(d)** genes.

**Fig. 4.**
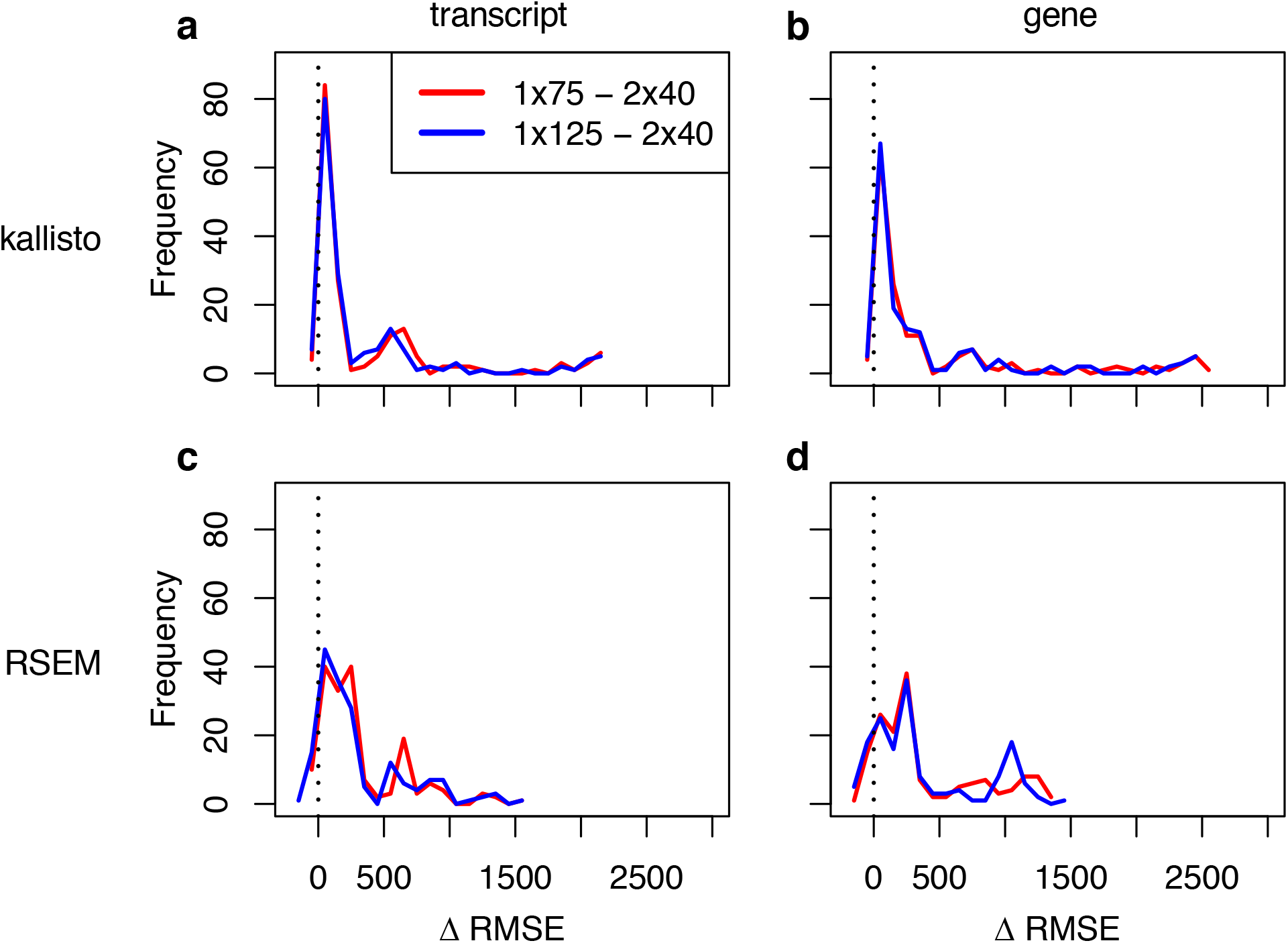
Histograms of differences in root-mean-square error (RMSE) between 2×40 and either 1×75 or 1×125, where RMSE is calculated using 2×125 as the gold standard. RMSE differences for kallisto for **(a)** transcripts and **(b)** genes, and RSEM for **(c)** transcripts and **(d)** genes. The vertical black dashed line indicates the threshold above which RMSE for either 1×75 or 1×125 is greater than that for 2×40.

We next evaluated the extent to which differences in performance with respect to estimating expression levels translated into downstream effects on tests of differential expression. Within each SRA accession and for each pair of conditions and for each sequencing strategy, we conducted Wald tests with, sleuth, limma-voom, and DESeq2. Similar to our evaluation of expression estimate robustness, we defined “true” differential expression signals as those recovered with 2×125, given a false discovery rate (FDR) of 0.01. We then calculated seven performance metrics on a per-accession basis for each differential expression method - sequencing strategy combination.

Sequencing strategy clearly impacts downstream differential expression analyses. Looking across all three differential expression methods, at both the transcript and gene level, for the majority of accessions false negative rates are lower for 2×40 compared to 1×75 (Figure 5; Additional file 4: Fig. S4, S5; Additional file 5: Table S2). The empirical false discovery rate— that is, for a strategy being evaluated, the proportion of putative differentially expressed features with a Benjamini-Hochberg adjusted p-value below our chosen FDR threshold of 0.01for which the adjusted p-value is >0.01 with 2×125—was also lower for 2×40 compared to both 1×75 and x125, although it was consistently higher than 0.01, the threshold used to classify Wald tests as significant, including the 2×125 analyses with which we defined true positives and negatives (Figure 5; Additional file 4: Fig. S4, S5; Additional file 5: Table S2). FDR control was poorer at the transcript than the gene level, and, as previously reported, DESeq2 did not control FDR as well as limma-voom and sleuth (18). At both the transcript and gene level, AUC is greater for 2×40 than 1×75 for all but a handful of method-accession combinations (Figure 5, Additional file 4: Fig. S4, S5; Additional file 5: Table S2). Additional metrics indicate a similar overall performance advantage of 2×40 compared to 1×75 (Additional file 4: Table S2). For all metrics, the short paired-end strategy outperforms 1×125 for the majority of method-accession combinations, although the magnitude of performance advantage of 2×40 compared to 1×125 is smaller than when compared with 1×75 smaller (Figure 5, Additional file 4: Fig. S4,S5; Additional file 5: Table S2).

**Fig. 5.**
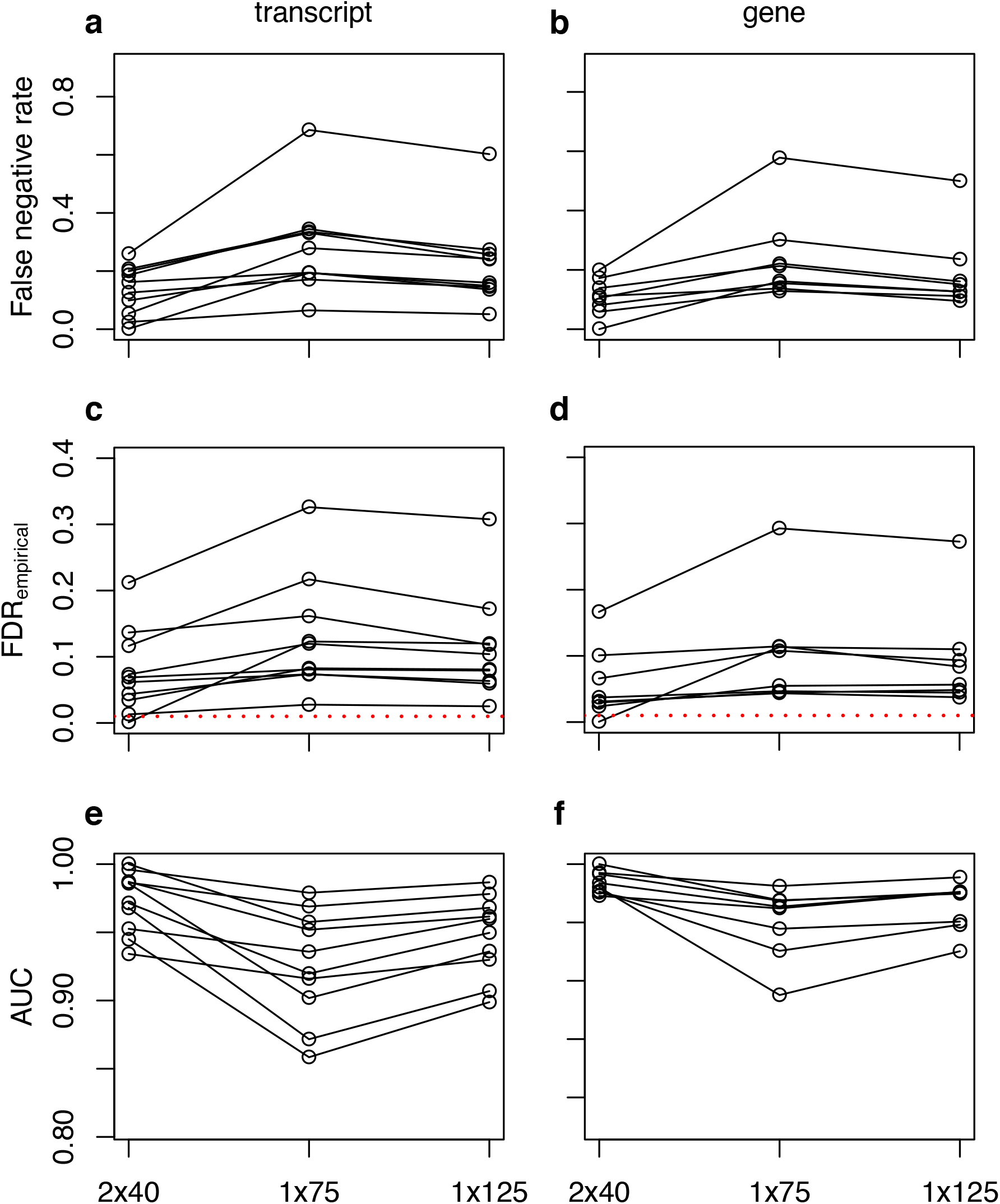
Variation across SRA accessions in **(a,b)** false negative rate, **(c,d)** empirical false discovery rate, and **(e,f)** AUC between 2×40, 1×75 and 1×125 sequencing strategies based upon differential expression testing with sleuth. Differences are plotted as means for Wald tests within accessions, for **(a,c,e)** transcripts and **(b,d,f)** genes.

## CONCLUSIONS

When designing a gene expression study using bulk RNA-seq, given a fixed budget, more robust expression estimates and differential expression test results can be obtained from short paired-end reads compared to longer single-end reads. In other words, the extra information provided by reads pairs during the mapping or pseudoalignment phase outweighs any potential penalty of short reads. In almost all cases, short paired-end reads trump single-end reads that sequence approximately the same number of bases. Short paired-end reads provide an additional benefit in that they can identify a large fraction of differentially expressed isoforms, for the cost of gene-level analysis with longer single-end reads. From a practical perspective, neither NextSeq nor NovaSeq sequencing kits distinguish between single and paired-end libraries. In the experience of support staff at the Bauer sequencing core facility at Harvard, one can get 92 bp from the NextSeq 75 bp kit, such that 2×40 is a viable sequencing design. Alternatively, the smallest NovaSeq kit is for 100 cycles, which can be used to sequence paired 50bp reads, allowing one to leverage that instrument’s much smaller cost per sequenced base. Overall, our findings suggest that for a given number of sequencing cycles, the most efficient design for RNA-seq expression analysis is sequencing shorter paired reads, rather than longer single-end reads.

## METHODS

### Datasets

From NCBI’s Short Read Archive, we downloaded paired-end RNA-seq reads from 12 different studies (Table 1), representing model organisms with well-annotated reference genomes: mouse (*Mus musculus*), human (*Homo sapiens*), yeast (*Saccharomyces cerevisiae*), fly (*Drosophila melanogaster*), and worm (*Caenorhabditis elegans*). In selecting data sets, we required minimum pre-processing read lengths to be ≥ 125 bp, and experimental designs to contain ≥ 2 experimental conditions.

### Short-read processing

We removed adapters and trimmed reads with NGmerge (21). For each sample, we trimmed reads to 125, 75, and 40bp using awk. For downstream analyses, we used 2×125, 2×40,1×125, and 1×75 trimmed data sets. Single-end data sets were comprised of the first read from the original paired reads. Reads were trimmed from the distal ends, such that trimming mimicked how sequencing to different lengths would be carried out on an Illumina sequencer.

### Expression analyses

For each sample-sequencing strategy combination, we estimated transcript-level expression with kallisto (22). We designated abundance estimates derived from 2×125 as our gold standard against which to benchmark all other strategies. We used Spearman’s rank correlations to quantify the degree of concordance between alternative sequencing strategies and the gold standard in terms of transcript per million (TPM). We performed similar analyses at the gene level by aggregating transcript-level TPMs using the R package Tximport (19). For yeast, which has no alternative splicing, analyses were only conducted at the transcript level. As an additional estimate of the correspondence between alternative sequencing strategies with 2×125 we also calculated root-mean-square error (RMSE).

Because a large proportion of RNA-seq applications involve tests of differential expression (DE), we compared differential expression test results between the gold standard 2×125 and alternative strategies. Specifically, for each experiment we conducted all possible pairwise Wald tests of transcript-level DE with sleuth (18) version 0.30.0, and used p-value aggregation (https://pachterlab.github.io/sleuth_walkthroughs/pval_agg/analysis.html) for gene-level analyses. To compare performance of alternative strategies relative to the gold standard, we calculated false negative and false positive rates, defining significant tests as those with Benjamini-Hochberg adjusted p-values ≤ 0.01.

In order to evaluate whether our findings were pipeline-specific, perhaps related to particular features of tools such as kallisto that rely on pseudo-alignment, we also performed expression analyses with an alternative pipeline, involving estimation of expression levels with RSEM (23) version 1.2.29 based upon alignments with bowtie2 version 2.3.2, (24). When SRA metadata enabled us to determine that libraries were stranded, we included the relevant RSEM command line arguments for restricting bowtie2 alignments to the correct read pair configuration. Using these expression estimates, we conducted differential expression tests with two different methods: limma-voom (25) version 3.38.3 and DESeq2 (26) version 1.22.2. For limma-voom, we only conducted tests where, for each gene or transcript,there were ≥ 2 samples with counts per million (CPM) ≥ 1. For limma-voom, we employed TMM normalization, and the voomWithQualityWeights option. Sleuth and limma-voom have previously been demonstrated to be top performers that correctly control FDR (22), and while DESeq2 has been shown to not control FDR as well as these two methods (22), we included it because it is a commonly used differential expression method. For all differential expression analyses, we calculated performance metrics for Wald tests where, at the specified false discovery rate (FDR) threshold of 0.01, there were ≥ 50 significantly differentially expressed features in all sequencing strategies. Metrics were calculated relative to 2×125, i.e. significant and non-significant results obtained with 2×125 were treated as true positives and negatives, respectively. Those metrics were the empirical false discovery rate, false positive rate, false negative rate, sensitivity, specificity, precision, and area under the curve (AUC) based upon receiver-operator statistics. These metrics were calculated with python scripts available at our associated github repository (see below). As the multiple-test adjusted p-values are the standard metric with which to determine whether particular transcripts or genes are differentially expressed, for AUC calculations we used these values as the response variable relative to the “true” classification of differences derived from tests based upon the 2×125 data. All reference genomes and gtf-formatted annotations against which to estimate expression were downloaded from Ensembl, and transcripts for kallisto-based analyses were extracted using the *extract-reference-transcripts* utility in RSEM.

## DECLARATIONS

### Availability of data and materials

The datasets analyzed during the current study and which support the conclusions of this article are publicly available Illumina short read data available on NCBI’s Short Read Archive (https://www.ncbi.nlm.nih.gov/sra). Specific accessions are listed in Additional File 1: Table S1. Command lines for executing NGmerge, kallisto, RSEM, sleuth, and limma-voom analyses are available on GitHub at https://github.com/harvardinformatics/rnaseq_readlength_assessment.

## Ethics approval and consent to participate

Not applicable

## Consent for publication

Not applicable

## Authors’ contributions

TBS and AHF designed the experiments, AHF and JMG analyzed the data, and AHF wrote the paper with TBS.

## Acknowledgments

The computations in this paper were run on the Odyssey cluster supported by the FAS Division of Science, Research Computing Group at Harvard University. We thank Claire Reardon, director of the Bauer Core at Harvard University for helpful discussions during the study design phase of our study, and for feedback on preliminary results. We thank Emma White at the Bauer Core for details concerning available sequencing read lengths and kit sizes for Illumina instruments. We also thank members from the Harvard FAS Informatics Group for feedback on preliminary results produced by this study. This work was conducted on the traditional territory of the Wampanoag and Massachusett peoples.

