## supplemental tables and figures for "Short paired-end reads trump long single-end reads for expression analysis"

**Table S1.** SRA accessions used in this study.

| Accession | Organism | Instrument | Paired-end read length | Conditions | Biological replicates per condition | Kallisto | RSEM |
|---|---|---|---|---|---|---|---|
| SRP133853 | *H. sapiens* | HiSeq 2500 | 125 | 6 | 3 | x | x |
| SRP115815 | *H. sapiens* | HiSeq 2500 | 125-126 | 9 | 3 | x | |
| SRP105271 | *M. musculus* | HiSeq 2000 | 125 | 5 | 3-4 | x | |
| SRP143508 | *M. musculus* | HiSeq X Ten | 150 | 2 | 3 | x | |
| SRP096374 | *M. musculus* | HiSeq 4000 | 150 | 2 | 3 | x | x |
| ERP017328 | *D. melanogaster* | HiSeq 2500 | 126 | 2 | 3 | x | |
| SRP128516 | *D. melanogaster* | HiSeq 4000 | 151 | 2 | 3 | x | x |
| SRP089981 | *C. elegans* | NextSeq 500 | 151 | 4 | 5 | x | |
| SRP092256 | *C. elegans* | HiSeq 2500 | 126 | 7 | 5 | x | |
| SRP129557 | *C. elegans* | HiSeq 3000 | 126 | 4 | 3 | x | x |
| SRP133093 | *S. cerevisiae* | HiSeq 2000 | 151 | 4 | 3 | x | x |
| SRP142501 | *S. cerevisiae* | HiSeq X Ten | 150 | 2 | 3 | x | |

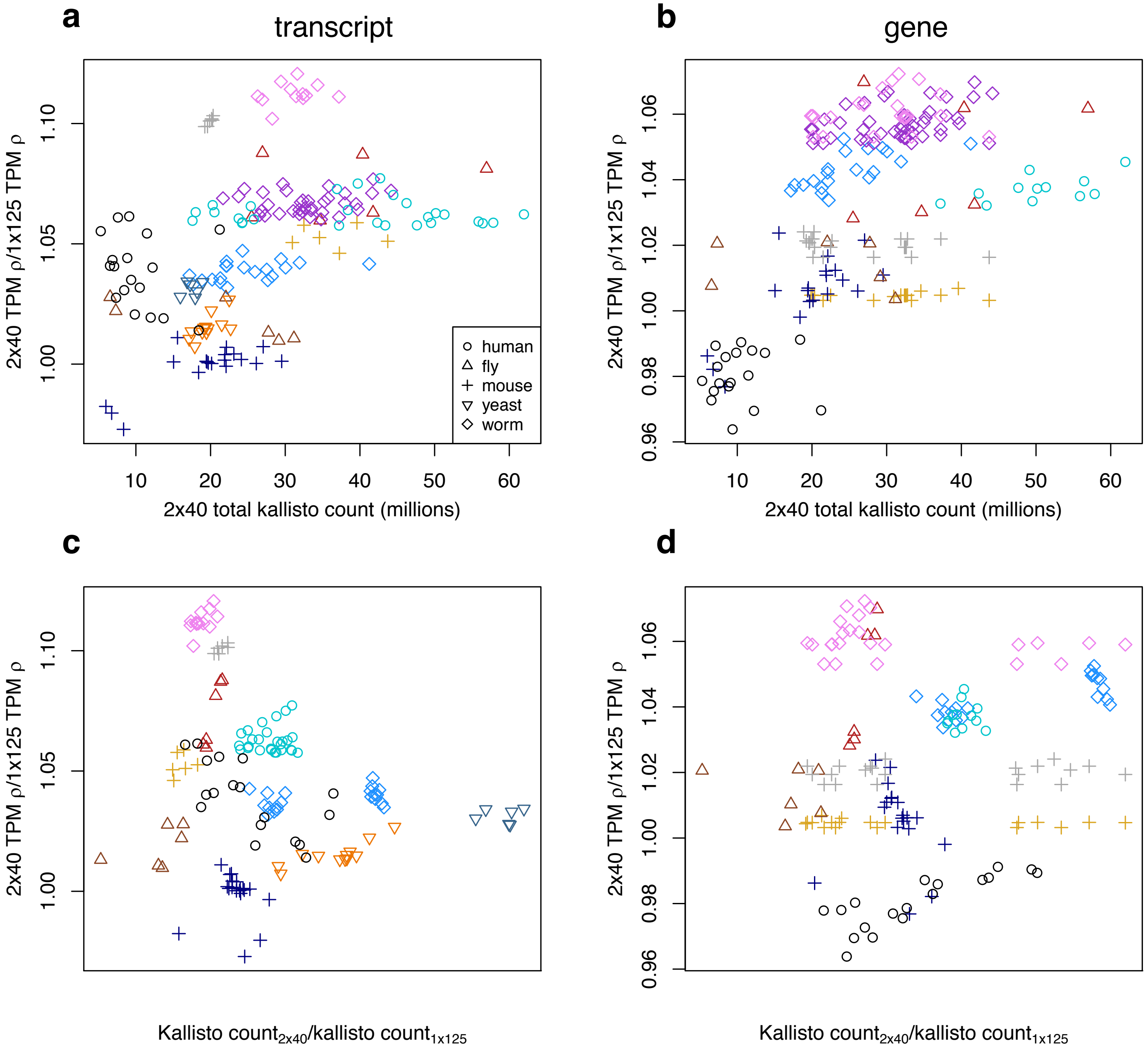

**Figure S1.** Effects of kallisto pseudo-alignment efficiency on differences in TPM estimate Spearman's rank correlations between 2x40 vs. 1x125. Efficiency of 2x40 is presented as the total kallisto count for 2x40 for **(a)** transcripts and **(b)** genes, and contrasted with the efficiency of 1x125 in the form of count ratios for **(c)** transcripts and **(d)** genes.

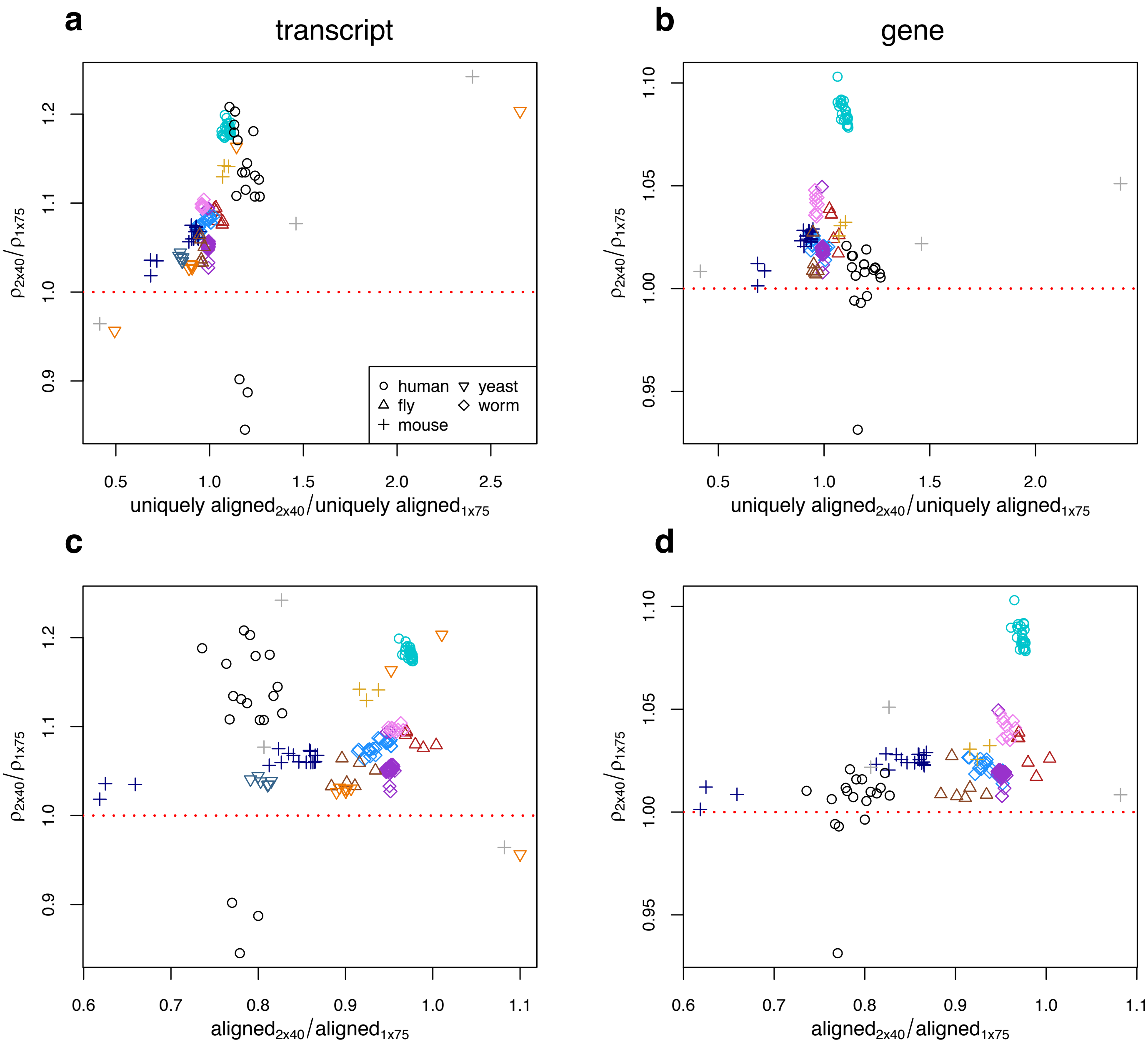

**Figure S2.** Effects of bowtie2 alignment rates on differences in RSEM TPM estimate Spearman's rank correlations between 2x40 vs. 1x75. Relative differences in efficiency are presented as the ratio of 2x40 to 1x75 TPM correlations with 2x125 TPM over the ratio of unique alignment rate for **(a)** transcripts and **(b)** genes, and over the ratio of overall alignment rates for **(c)** transcripts and **(d)** genes.

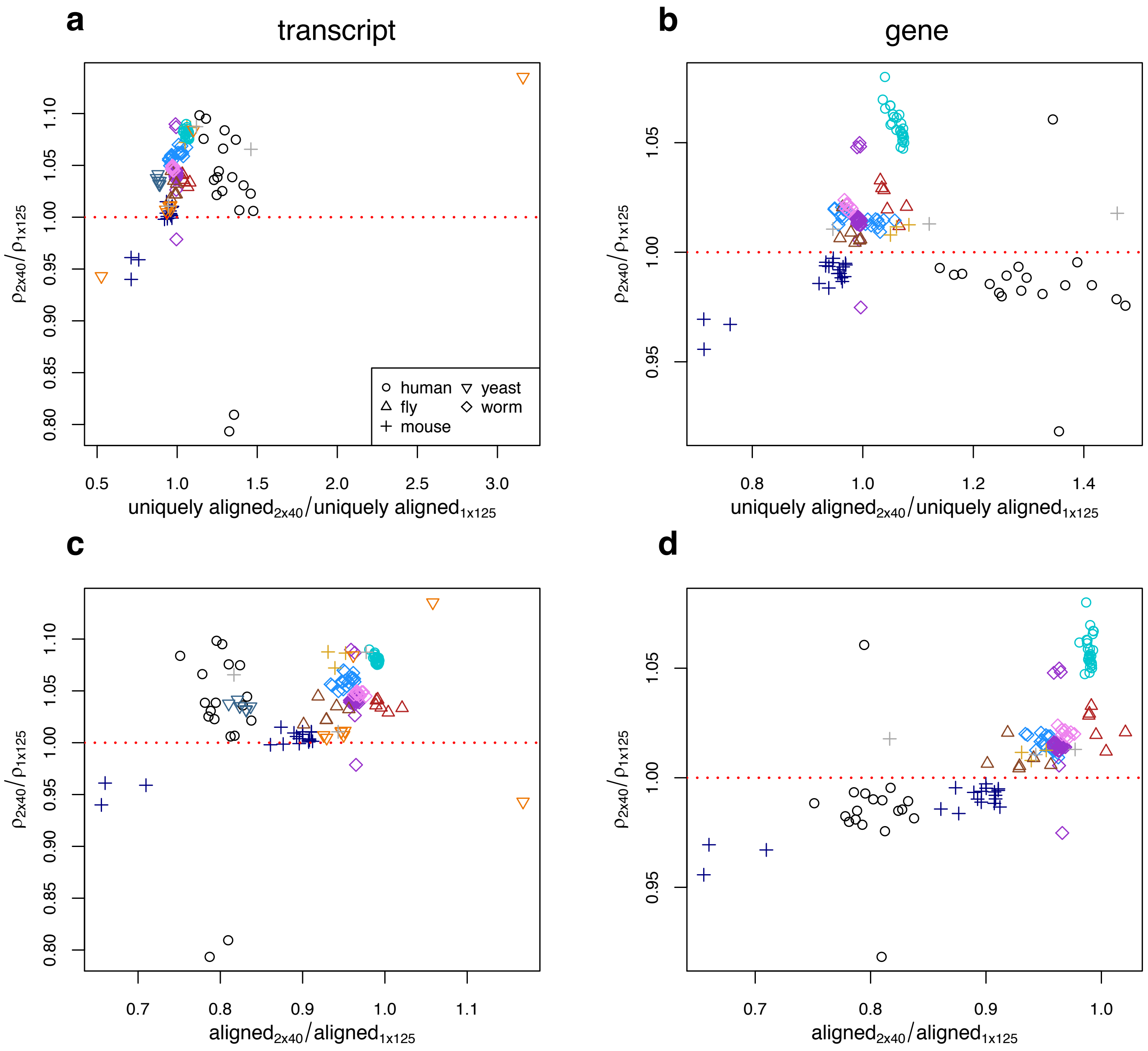

**Figure S3.** Effects of bowtie2 alignment rates on differences in RSEM TPM estimate Spearman's rank correlations between 2x40 vs. 1x125. Relative differences in efficiency are presented as the ratio of 2x40 to 1x125 TPM correlations with 2x125 TPM over the ratio of unique alignment rate for **(a)** transcripts and **(b)** genes, and over the ratio of overall alignment rates for **(c)** transcripts and **(d)** genes.

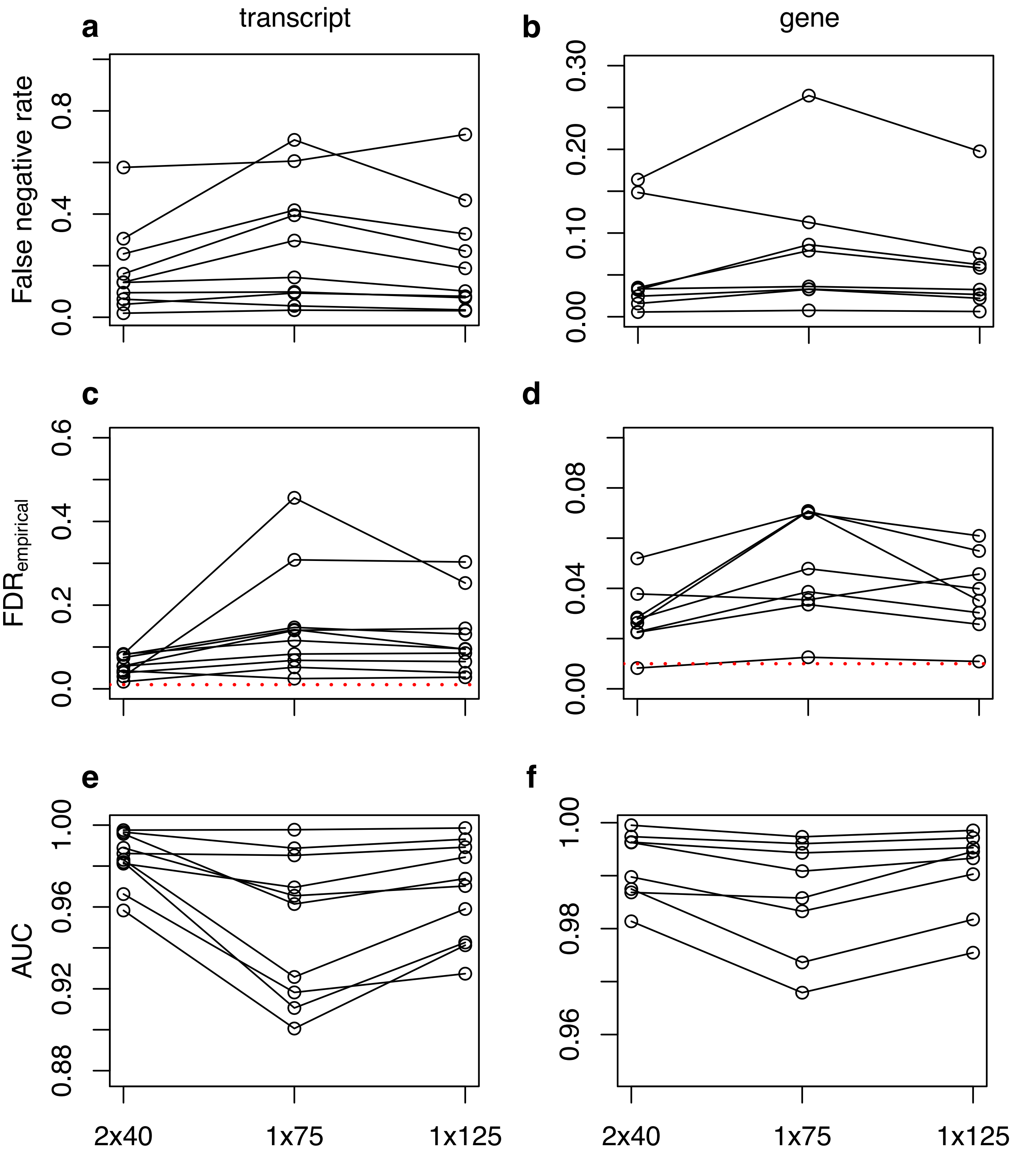

**Figure S4.** For limma-voom Wald tests, variation across SRA accessions, in **(a,b)** false negative rate**, (c,d)** empirical false discovery rate (red line indicates FDR threshold of 0.01 for calling tests significant), and **(e,f)** AUC between 2x40, 1x75 and 1x125 sequencing strategies. Differences are plotted as means for Wald tests within accessions, for **(a,c,e)** transcripts and **(b,d,f)** genes.

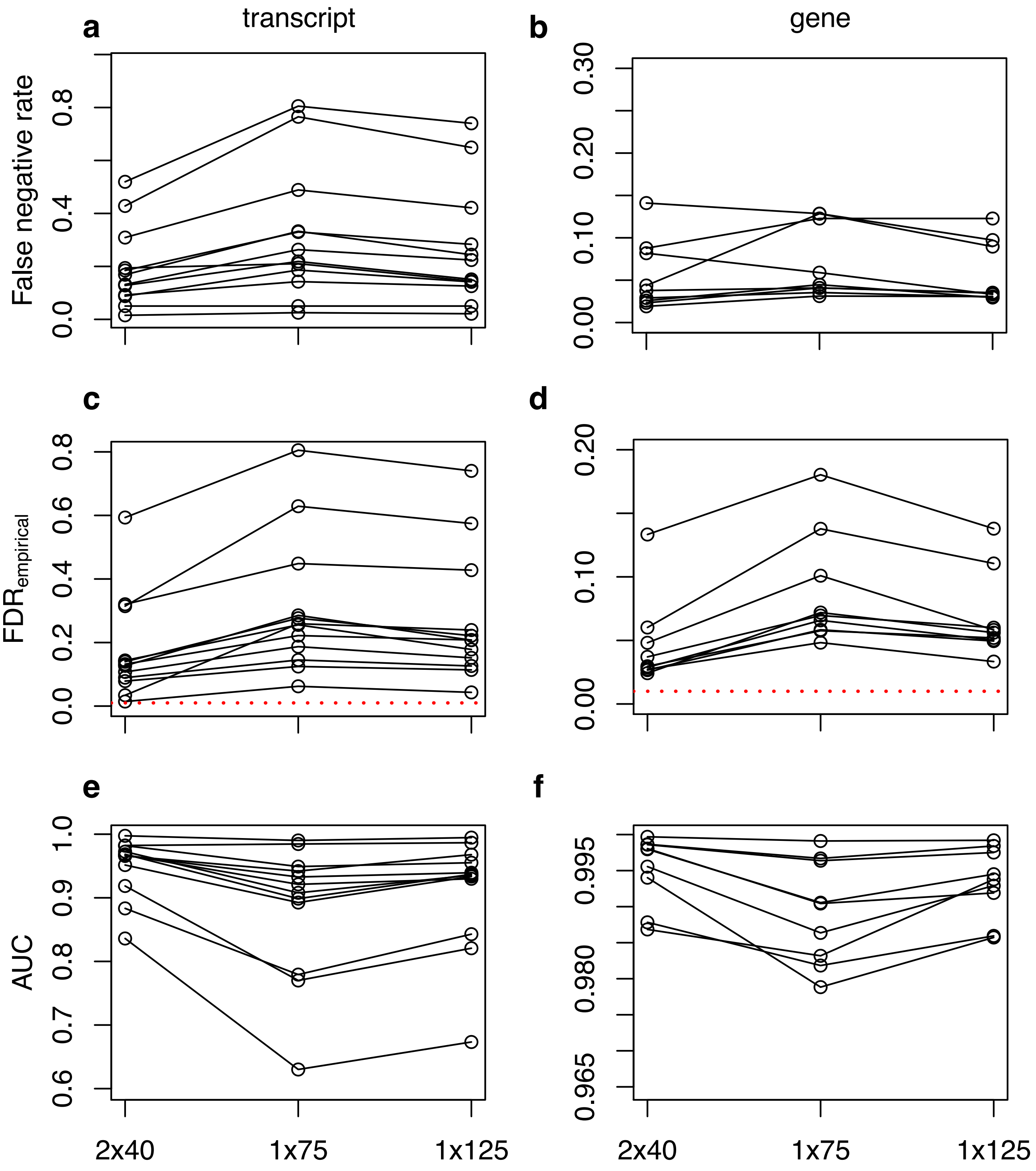

**Figure S5.** For DESeq2 Wald tests, variation across SRA accessions, in **(a,b)** false negative rate**, (c,d)** empirical false discovery rate (red line indicates FDR threshold of 0.01 for calling tests significant), and **(e,f)** AUC between 2x40, 1x75 and 1x125 sequencing strategies. Differences are plotted as means for Wald tests within accessions, for **(a,c,e)** transcripts and **(b,d,f)** genes.

**Table S2**. Across all assayed SRA accessions, percentages of pairwise differential expression tests between conditions where a performance metric is greater for 2x40 compared to 1x75 or 1x125. Gene-level analyses are not included for yeast accessions (SRP133093 and SRP142501) as this species has no alternative splicing, such that there is only one annotated transcript per gene. Bold values indicate those where 2x40 outperforms the evaluated single-end strategy. NA indicates there were < 50 differentially expressed features such that performance metrics were not calculated.

| | transcript | | gene | |
|---|---|---|---|---|
| **Metric** | **2x40 > 1x75** | **2x40 > 1x125** | **2x40 > 1x75** | **2x40 > 1x125** |
| **sleuth** | | | | |
| **empirical false discovery rate** | | | | |
| ERP017328 | **0** | **0** | **0** | **0** |
| SRP089981 | **0** | 80 | **0** | **0** |
| SRP092256 | **0** | **0** | **0** | **0** |
| SRP096374 | **0** | **0** | **0** | **0** |
| SRP105271 | **0** | 90 | **0** | **0** |
| SRP115815 | **0** | **0** | **0** | **11.1** |
| SRP129557 | **0** | **0** | **0** | **0** |
| SRP133093 | **0** | **0** | **0** | **0** |
| SRP133853 | **46.7** | **46.7** | **46.7** | **46.7** |
| SRP142501 | **0** | **0** | **0** | **0** |
| **false negative rate** | | | | |
| ERP017328 | **0** | **0** | **0** | **0** |
| SRP089981 | **0** | **20** | **0** | **0** |
| SRP092256 | **0** | **0** | **0** | **0** |
| SRP096374 | **0** | **0** | **0** | **0** |
| SRP105271 | **0** | 100 | **0** | 100 |
| SRP115815 | **0** | **8.3** | **0** | **27.8** |
| SRP129557 | **0** | **0** | **0** | **0** |
| SRP133093 | **0** | **0** | – | – |
| SRP133853 | **0** | **0** | **13.3** | **13.3** |
| SRP142501 | **0** | **0** | – | – |
| **false positive rate** | | | | |
| ERP017328 | **0** | **0** | **0** | **0** |
| SRP089981 | 80 | 80 | 60 | 80 |
| SRP092256 | **0** | **0** | **0** | **0** |
| SRP096374 | **0** | **0** | **0** | **0** |
| SRP105271 | **0** | 90 | **20** | 80 |
| SRP115815 | **0** | **0** | **5.6** | **16.7** |
| SRP129557 | **0** | **0** | **0** | **0** |
| SRP133093 | **0** | **0** | – | – |
| SRP133853 | 60 | 60 | 53.3 | 53.3 |
| SRP142501 | **0** | **0** | – | – |

| | | | | |
|---|---|---|---|---|
| **sensitivity** | | | | |
| ERP017328 | **100** | **100** | **100** | **100** |
| SRP089981 | **100** | **80** | **100** | **100** |
| SRP092256 | **100** | **100** | **100** | **100** |
| SRP096374 | **100** | **100** | **100** | **100** |
| SRP105271 | **100** | 0 | **100** | 0 |
| SRP115815 | **100** | **91.7** | **100** | **72.2** |
| SRP129557 | **100** | **100** | **100** | **100** |
| SRP133093 | **100** | **100** | – | – |
| SRP133853 | **100** | **93.3** | **86.7** | **86.7** |
| SRP142501 | **100** | **100** | – | – |
| **specificity** | | | | |
| ERP017328 | **100** | **100** | **100** | **100** |
| SRP089981 | 20 | 20 | 40 | 20 |
| SRP092256 | **100** | **100** | **100** | **100** |
| SRP096374 | 0 | **100** | **100** | **100** |
| SRP105271 | **100** | 10 | **80** | 20 |
| SRP115815 | **100** | **100** | **94.4** | **83.3** |
| SRP129557 | **100** | **100** | **100** | **100** |
| SRP133093 | **100** | **100** | – | – |
| SRP133853 | 40 | 40 | 46.7 | **46.7** |
| SRP142501 | **100** | **100** | – | – |
| **precision** | | | | |
| ERP017328 | **100** | **100** | **100** | **100** |
| SRP089981 | 40 | 20.0 | **100** | 40 |
| SRP092256 | **100** | **100** | **100** | **100** |
| SRP096374 | **100** | **100** | **100** | **100** |
| SRP105271 | **100** | 10 | **100** | 20 |
| SRP115815 | **100** | **100** | **100** | **88.9** |
| SRP129557 | **100** | **100** | **100** | **100** |
| SRP133093 | **100** | **100** | – | – |
| SRP133853 | **53.3** | **53.3** | **53.3** | **53.3** |
| SRP142501 | **100** | **100** | – | – |
| **auc** | | | | |
| ERP017328 | **100** | **100** | **100** | **100** |
| SRP089981 | **100** | **80** | **100** | **100** |
| SRP092256 | **100** | **100** | **100** | **100** |
| SRP096374 | **100** | **100** | **100** | **100** |
| SRP105271 | **100** | 0 | **100** | **30** |
| SRP115815 | **100** | **100** | **100** | **100** |
| SRP129557 | **100** | **100** | **100** | **100** |
| SRP133093 | **100** | **100** | – | – |

| | | | | |
|---|---|---|---|---|
| SRP133853 | **100** | **100** | **100** | **100** |
| SRP142501 | **100** | **100** | – | – |
| **limma-voom** | | | | |
| **empirical false discovery rate** | | | | |
| ERP017328 | **0** | **0** | **0** | **0** |
| SRP089981 | **0** | **0** | **0** | **0** |
| SRP092256 | **0** | **0** | **0** | **5** |
| SRP096374 | 100 | 100 | 100 | **0** |
| SRP105271 | **0** | **0** | **0** | **40** |
| SRP115815 | **0** | **0** | **5.6** | **25** |
| SRP129557 | **0** | **0** | **0** | **0** |
| SRP133093 | **0** | **0** | **0** | **0** |
| SRP133853 | **25** | **25** | **42.9** | 50 |
| SRP142501 | **0** | **0** | **0** | **0** |
| **false negative rate** | | | | |
| ERP017328 (1) | **0** | **0** | **0** | **0** |
| SRP089981 (4) | **0** | **0** | **0** | **25** |
| SRP092256 (19) | **21** | 89.5 | **0** | **30** |
| SRP096374 (1) | **0** | **0** | **0** | **0** |
| SRP105271 (9) | **22.2** | 77.8 | 70 | 100 |
| SRP115815 (36) | **0** | **0** | **0** | **0** |
| SRP129557 (3) | **0** | **33.3** | **16.7** | **33.3** |
| SRP133093 (6) | **33.3** | 50 | – | – |
| SRP133853 ()??? | 50 | **25** | **35.7** | **42.9** |
| SRP142501 (1) | 100 | 100 | – | – |
| **false positive rate** | | | | |
| ERP017328 | **0** | **0** | **0** | **0** |
| SRP089981 | **0** | **0** | **0** | **0** |
| SRP092256 | **0** | **0** | **0** | **5.0** |
| SRP096374 | 100 | 100 | 100 | **0** |
| SRP105271 | **0** | **0** | **0** | **20** |
| SRP115815 | **11.1** | **0** | **8.3** | **30.6** |
| SRP129557 | **0** | **0** | **0** | **0** |
| SRP133093 | **0** | **0** | – | – |
| SRP133853 | 50 | 75 | 57.1 | 50 |
| SRP142501 | **0** | **0** | – | – |
| **sensitivity** | | | | |
| ERP017328 | **100** | **100** | **100** | **100** |
| SRP089981 | **100** | **100** | **100** | **75** |
| SRP092256 | **78.9** | 10.5 | **100** | **60** |
| SRP096374 | **100** | **100** | **100** | **100** |
| SRP105271 | **77.8** | 11.1 | 30 | 0 |

| | | | | |
|---|---|---|---|---|
| SRP115815 | **100** | **100** | **100** | **100** |
| SRP129557 | **100** | **66.7** | **83.3** | **66.7** |
| SRP133093 | **66.7** | 50 | – | – |
| SRP133853 | 50 | **75** | **64.3** | **57.1** |
| SRP142501 | 0 | 0 | – | – |
| **specificity** | | | | |
| ERP017328 | **100** | **100** | **100** | **100** |
| SRP089981 | **100** | **100** | **100** | **100** |
| SRP092256 | **100** | **100** | **100** | **95** |
| SRP096374 | 0 | 0 | 0 | **100** |
| SRP105271 | **100** | **100** | **100** | **80** |
| SRP115815 | **88.9** | **100** | **91.7** | **69.4** |
| SRP129557 | **100** | **100** | **100** | **100** |
| SRP133093 | **100** | **100** | – | – |
| SRP133853 | 50 | 25 | 42.9 | 50 |
| SRP142501 | **100** | **100** | – | – |
| **precision** | | | | |
| ERP017328 | **100** | **100** | **100** | **100** |
| SRP089981 | **100** | **100** | **100** | **100** |
| SRP092256 | **100** | **100** | **100** | **95** |
| SRP096374 | 0 | 0 | 0 | **100** |
| SRP105271 | **100** | **100** | **100** | **60** |
| SRP115815 | **100** | **100** | **94.4** | **75** |
| SRP129557 | **100** | **100** | **100** | **100** |
| SRP133093 | **100** | **100** | – | – |
| SRP133853 | **75** | **75** | **57.1** | 50 |
| SRP142501 | **100** | **100** | – | – |
| **auc** | | | | |
| ERP017328 | **100** | **100** | **100** | 0 |
| SRP089981 | **100** | **100** | **100** | **100** |
| SRP092256 | **100** | **100** | **100** | **75** |
| SRP096374 | 0 | 0 | **100** | **100** |
| SRP105271 | **100** | 33.3 | **60** | 10 |
| SRP115815 | **100** | **100** | **100** | **94.4** |
| SRP129557 | **100** | **100** | **100** | **100** |
| SRP133093 | **100** | **100** | – | – |
| SRP133853 | **100** | **100** | **100** | **71.4** |
| SRP142501 | **100** | 0 | – | – |
| **DESeq2** | | | | |
| **false discovery rate** | | | | |
| ERP017328 | **0** | **0** | **0** | **0** |
| SRP089981 | **0** | **0** | **0** | **20** |

| | | | | |
|---|---|---|---|---|
| SRP092256 | **4.8** | **9.5** | **0** | **4.8** |
| SRP096374 | **0** | **0** | **0** | **0** |
| SRP105271 | **0** | **0** | **0** | **20** |
| SRP115815 | **0** | **0** | **2.8** | **8.3** |
| SRP129557 | **0** | **0** | **0** | **0** |
| SRP133093 | **0** | **0** | **0** | **0** |
| SRP133853 | **0** | **0** | **13.3** | **20** |
| SRP142501 | **0** | **0** | **0** | **0** |
| SRP128516 | **0** | **0** | **0** | **0** |
| SRP143508 | **0** | **0** | **0** | **0** |
| **false negative rate** | | | | |
| ERP017328 | **0** | **0** | **0** | **0** |
| SRP089981 | **0** | **0** | **0** | **0** |
| SRP092256 | **0** | **0** | **9.5** | **42.9** |
| SRP096374 | **0** | **0** | **0** | 100 |
| SRP105271 | **20** | 100 | 90 | 100 |
| SRP115815 | **0** | **0** | **0** | **0** |
| SRP129557 | **0** | **0** | **0** | **16.7** |
| SRP133093 | **0** | **33.3** | – | – |
| SRP133853 | **0** | **20** | 66.7 | 93.3 |
| SRP142501 | **0** | **0** | – | – |
| SRP128516 | **0** | **0** | **0** | **0** |
| SRP143508 | **0** | **0** | NA | NA |
| **false positive rate** | | | | |
| ERP017328 | **0** | **0** | **0** | **0** |
| SRP089981 | **0** | **0** | **0** | **20** |
| SRP092256 | **9.5** | **14.3** | **0** | **4.8** |
| SRP096374 | **0** | **0** | **0** | **0** |
| SRP105271 | **0** | **0** | **0** | **20** |
| SRP115815 | **0** | **0** | **8.3** | **11.1** |
| SRP129557 | **0** | **16.7** | **0** | **0** |
| SRP133093 | **0** | **0** | – | – |
| SRP133853 | **46.7** | **40** | **20** | **20** |
| SRP142501 | **0** | **0** | – | – |
| SRP128516 | **0** | **0** | **0** | **0** |
| SRP143508 | **0** | **0** | NA | NA |
| **sensitivity** | | | | |
| ERP017328 | **100** | **100** | **100** | **100** |
| SRP089981 | **100** | **100** | **100** | **100** |
| SRP092256 | **100** | **100** | **90.5** | **52.4** |
| SRP096374 | **100** | **100** | **100** | 0 |
| SRP105271 | **80** | 0 | 10 | 0 |

| | | | | |
|---|---|---|---|---|
| SRP115815 | **100** | **100** | **100** | **100** |
| SRP129557 | **100** | **100** | **100** | **66.7** |
| SRP133093 | **100** | **66.7** | – | – |
| SRP133853 | **100** | **80** | 33.3 | 6.7 |
| SRP142501 | 0 | 0 | – | – |
| SRP128516 | **100** | **100** | **100** | **100** |
| SRP143508 | **100** | **100** | NA | NA |
| **specificity** | | | | |
| ERP017328 | **100** | **100** | **100** | **100** |
| SRP089981 | **80** | **100** | **100** | **80** |
| SRP092256 | **90.5** | **85.7** | **100** | **95.2** |
| SRP096374 | **100** | **100** | **100** | **100** |
| SRP105271 | **100** | **100** | **100** | **80** |
| SRP115815 | **100** | **100** | **91.7** | **88.9** |
| SRP129557 | **100** | **83.3** | **100** | **100** |
| SRP133093 | **100** | **100** | – | – |
| SRP133853 | **53.3** | **60** | **80** | **80** |
| SRP142501 | **100** | **100** | – | – |
| SRP128516 | **100** | **100** | **100** | 0 |
| SRP143508 | **100** | **100** | NA | NA |
| **precision** | | | | |
| ERP017328 | **100** | **100** | **100** | **100** |
| SRP089981 | **100** | **100** | **100** | **80** |
| SRP092256 | **95.2** | **90.5** | **100** | **95.2** |
| SRP096374 | **100** | **100** | **100** | **100** |
| SRP105271 | **100** | **100** | **100** | **80** |
| SRP115815 | **100** | **100** | **97.2** | **91.7** |
| SRP129557 | **100** | **100** | **100** | **100** |
| SRP133093 | **100** | **100** | – | – |
| SRP133853 | **100** | **100** | **86.7** | **80** |
| SRP142501 | **100** | **100** | – | – |
| SRP128516 | **100** | **100** | **100** | **100** |
| SRP143508 | **100** | **100** | NA | NA |
| **auc** | | | | |
| ERP017328 | **100** | **100** | **100** | **100** |
| SRP089981 | **100** | **100** | **100** | **100** |
| SRP092256 | **100** | **100** | **100** | **100** |
| SRP096374 | **100** | **100** | **100** | **100** |
| SRP105271 | **100** | 40 | **90** | 0 |
| SRP115815 | **100** | **100** | **100** | **100** |
| SRP129557 | **100** | **100** | **100** | **100** |
| SRP133093 | **100** | **100** | – | – |

| | | | | |
|---|---|---|---|---|
| SRP133853 | **100** | **100** | **93.3** | **73.3** |
| SRP142501 | 0 | 0 | – | – |
| SRP128516 | **100** | **100** | **100** | **100** |
| SRP143508 | **100** | **100** | NA | NA |
